# DeleteomeTools: Utilizing a compendium of yeast deletion strain transcriptomes to identify co-functional genes

**DOI:** 10.1101/2024.02.05.578946

**Authors:** Maxwell L. Neal, Sanjeev K. Choudhry, John D. Aitchison

## Abstract

We introduce DeleteomeTools, an R package that leverages the Deleteome compendium of yeast single-gene deletion transcriptomes to predict gene function. Primarily, the package provides functions for identifying similarities between the transcriptomic signatures of deletion strains, thereby associating genes of interest with others that may be functionally related. We describe how our software predicted a novel relationship between the yeast nucleoporin Nup170 and the Ctf18-RFC complex, which was confirmed experimentally, revealing a previously unknown link between nuclear pore complexes and the DNA replication machinery. To assess the package’s broader predictive capabilities, we performed a systematic evaluation that tested how well it predicted Gene Ontology (GO) annotations already applied to the subset of genes deleted in Deleteome strains. We show that our package predicted a majority of reported GO:*biological process* annotations with semantic similarities ranging from moderate to identical. We also discuss how our strategy for quantifying similarity between deletion strains, which relies on differential expression signatures, differs from other approaches that use global expression profiles and why it has the potential to identify functional relationships that might otherwise go undetected.

## INTRODUCTION

A central motivation for systems biology research is to understand the function of complex biological systems from a holistic perspective. Within the context of a single cell, systems research can illuminate how the interplay of thousands of gene products produces a cellular phenotype of interest. The increasing availability of public, omics-level data resources provide holistic readouts from biological systems serving as a valuable foundation to address this challenge. One such resource is the Deleteome compendium (1) which comprises 1,484 transcriptomic profiles obtained from *Saccharomyces cerevisiae* single-gene deletion strains. The Deleteome’s creators used microarrays to measure transcript abundances for each deletion strain, comparing them to wild-type controls. The compendium includes expression changes and associated *P*-values derived from comparisons, providing a transcriptomic “signature” for each deletion strain based on its set of differentially expressed genes. The Deleteome has been applied to generate and analyze systems-level gene regulatory networks, identify gene-specific transcriptional repressors, and predict novel protein-protein interactions in yeast (1, 2).

We used the Deleteome to functionally characterize the nucleoporin Nup170, which is a structural constituent of the nuclear pore complex (NPC) and also plays a role in gene regulation, including sub-telomeric silencing (3, 4). In that study, to help reveal the mechanisms whereby Nup170 mediates silencing, we designed and implemented R code for collecting the differential expression signature of the Nup170 deletion strain (*nup170*Δ) in the Deleteome, iteratively comparing it to signatures from all other strains in the compendium, and then identifying and ranking deletion strains with similar signatures (4). We designed our codebase so that this task of identifying transcriptionally similar deletion strains could be performed for any strain in the compendium, not just for *nup170*Δ. These similarity analyses were critical to our discovery of Nup170’s functional association with the chromosome transmission fidelity 18-replication factor C (Ctf18-RFC) complex (4). Among the transcriptome signatures similar to *nup170*Δ were transcriptomes from chromosome transmission fidelity 18 (*ctf18*Δ), chromosome transmission fidelity 8 (*ctf8*Δ), and defective in sister chromatid cohesion 1 (*dcc1*Δ) strains. These represent the three non-essential components of the Ctf18-RFC complex, which helps establish chromosome cohesion during mitosis and is involved in loading and unloading proliferating cell nuclear antigen (PCNA). (Deletion strains for the other members of the Ctf18-RFC complex, due to their essentiality, are not present in the Deleteome and could not have been identified as co-functional with Nup170 using our analysis.) Our subsequent experimental investigations demonstrated that the Ctf18-RFC complex is recruited to a subset of distinctly-structured NPCs, revealing a novel association between NPCs, the Ctf18-RFC complex, and PCNA, which is required for DNA replication and repair (4). Given that our computational analyses predicted this co-functional relationship between Nup170 and the Ctf18-RFC complex, we have made our code publicly available, and validated the approach with other strains, in the hopes that the research community may find it useful for identifying novel functional relationships among other genes.

In the following sections, we detail the computational methods implemented in DeleteomeTools for quantifying similarity between deletion strain transcriptomes and predicting gene function as well as additional visualization and analysis features that it provides. We also report results from a systematic evaluation where we assessed our tool’s capabilities for predicting gene function, and which identified optimal statistical significance thresholds to use when making such predictions.

## MATERIAL AND METHODS

DeleteomeTools was developed to quantify the similarity between transcriptomic abundances in a deletion strain of interest, which we term a “query strain” containing a deleted “query gene”, and all other Deleteome strains. In a broad sense, the software performs phenotype matching: the set of gene expression values present in the query strain constitutes a phenotype that is compared against expression values in other strains. Then, functionally profiling the genes deleted in matching strains can predict the functionality of the gene deleted in the query strain. This approach for predicting gene function from a compendium of single-gene deletion strains has been applied previously using a smaller set of strains (5), and we aimed to apply a similar approach using the more comprehensive Deleteome dataset and updated functional annotations. Figure 1 summarizes our methodology for gene function prediction and illustrates the main features of the DeleteomeTools package.

**Figure 1.**
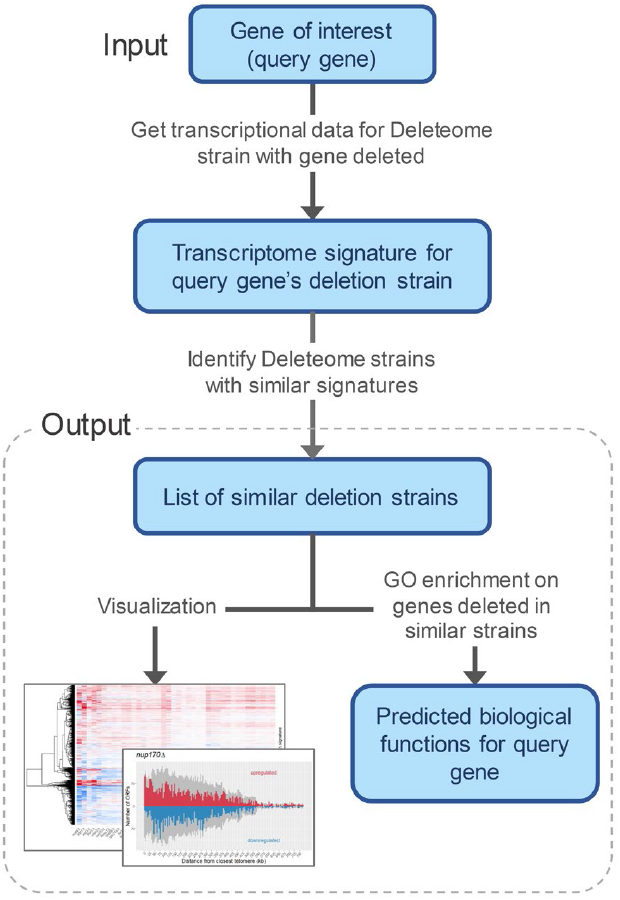
Main steps of the DeleteomeTools pipeline wherein a compendium of deletion strain transcriptomes is used to predict biological functions for a gene of interest.

After DeleteomeTools computes similarity scores between a query strain and all other Deleteome strains, scores are ranked and thresholds applied to remove lower-confidence scores, thereby generating a list of deletion strains similar to the query strain. The software provides two complementary methods for quantifying strain similarity as well as visualization and analysis features, which we detail below.

### Strain similarity based on reciprocal correlation

The first similarity metric provided by DeleteomeTools is based on correlation tests. In the following example, we detail how this metric is computed given a query strain of interest (“Strain A”) and another strain from the Deleteome to which it is being compared (“Strain B”). First, the set of differentially expressed genes in Strain A (those significantly different from wild-type based on *P*-values provided in the Deleteome) are identified. These genes constitute Strain A’s transcriptomic “signature”. We then compare the expression values of signature genes from Strain A to the corresponding values in Strain B. If the correlation is statistically significant and positive, we perform a reciprocal test where expression values of Strain B’s signature genes are correlated against those in strain A. This step helps ensure that the complete signature of Strain A is compared to the complete signature of Strain B, as additional components in one signature could indicate lower similarity between the strains. Consider the case where the function of the gene deleted in Strain A is limited to a specific regulatory pathway and the gene deleted in Strain B functions in that pathway but also in several others. A correlation test between Strains A and B limited to the differentially-expressed genes in Strain A’s deletion signature might show significance but would ignore the larger changes present in Strain B’s signature. In this example, the reciprocal correlation test may not prove significant since deletion of the gene in Strain B would introduce a larger number of transcriptional changes which might not be present in Strain A. Thus, if a reciprocal correlation test is insignificant or negative, the two strains are not classified as similar. Because the gene deleted in Strain B has additional functionality compared to the gene deleted in Strain A, our tools may not necessarily classify the genes as similar, even though their functions may partly overlap.

DeleteomeTools performs this reciprocal correlation test between a query strain and all other deletion strains in the Deleteome and then collects the strains that were significantly, positively correlated in both directions. It then ranks strain similarity based on FDR-corrected *P*-values from the first correlation test (expression values of Strain A’s signature compared to corresponding values in Strain B). In initial tests using this approach, we found that some deletion mutants, including *nup170*Δ, had several hundred similar deletion strains. We therefore added a feature to down-select this list by including only those strains whose correlation *P*-values are within a certain percentile of all *P*-values. Given the strong correlation between these *P*-values and their corresponding correlation coefficients, this down-selection serves to filter out statistically-significant correlations with low effect sizes. In our work with *nup170*Δ, we used the 95^th^ percentile as our similarity cutoff, although users can readily adjust thresholds for gene expression values and FDR-corrected correlation test *P*-values used in analyses.

### Strain similarity based on signature enrichment

The second strain similarity metric implemented in DeleteomeTools uses a more qualitative approach compared to reciprocal correlation. We developed this alternative because reciprocal correlation tests may not always be significant for co-functional genes, given that they rely on quantitative gene expression values. Consider an example involving two Deleteome Strains A and B where the gene deleted in A co-functions in a transcriptional regulatory pathway with the gene deleted in B. The specific expression changes introduced in Strain A may not show a significant correlation with those in B, even though the expression changes are qualitatively similar: expression of all downstream genes in the pathway might increase in both strains, but the specific expression values might not correlate significantly. Therefore, we developed an “enrichment-for-signature” similarity test that could account for such cases. In this approach, the genes in the signature of a deletion strain of interest are identified using the same approach described above, and the expression of each gene is categorized as either increased or decreased based on its value. Then, when comparing that strain’s signature to that of another, we collect the signature of the second strain and label its specific genes as increased or decreased, and then identify the genes that are present in both signatures and have the same directional expression change (i.e., genes that increased significantly in both strains or decreased significantly in both). We then perform a hypergeometric test to quantify the probability of observing that number of overlapping genes between the two signatures by chance. After performing hypergeometric tests comparing a query strain to all other Deleteome strains, we FDR-correct the associated *P*-values, select those that are significant, and apply a percentile cutoff as described above to determine which strains qualify as similar.

In our experience, the enrichment-based approach tends to be less conservative than the reciprocal correlation option as it often identifies a comparatively higher number of similar strains. However, we encourage the complementary use of both approaches, as they are non-redundant. In our original analyses on *nup170*Δ, the enrichment-based approach identified 74 similar strains and the correlation-based approach identified 39. Twenty-seven were common to both (including strains for all three components of the Ctf18-RFC complex), indicating that there is likely to be substantial overlap in the results from each method, but they may also be complementary.

### Visualization features

DeleteomeTools also provides functions for visualizing transcriptome data across multiple deletion strains and the relative genomic position of a deletion strain’s differentially expressed genes. For the former, users can easily generate gene expression heatmaps for a given deletion strain alongside any other strains in the compendium. An example is shown in Figure 2 where the expression values for genes in the *nup170*Δ strain are shown in addition to those from the 39 similar deletion strains identified using the reciprocal correlation method described above.

**Figure 2.**
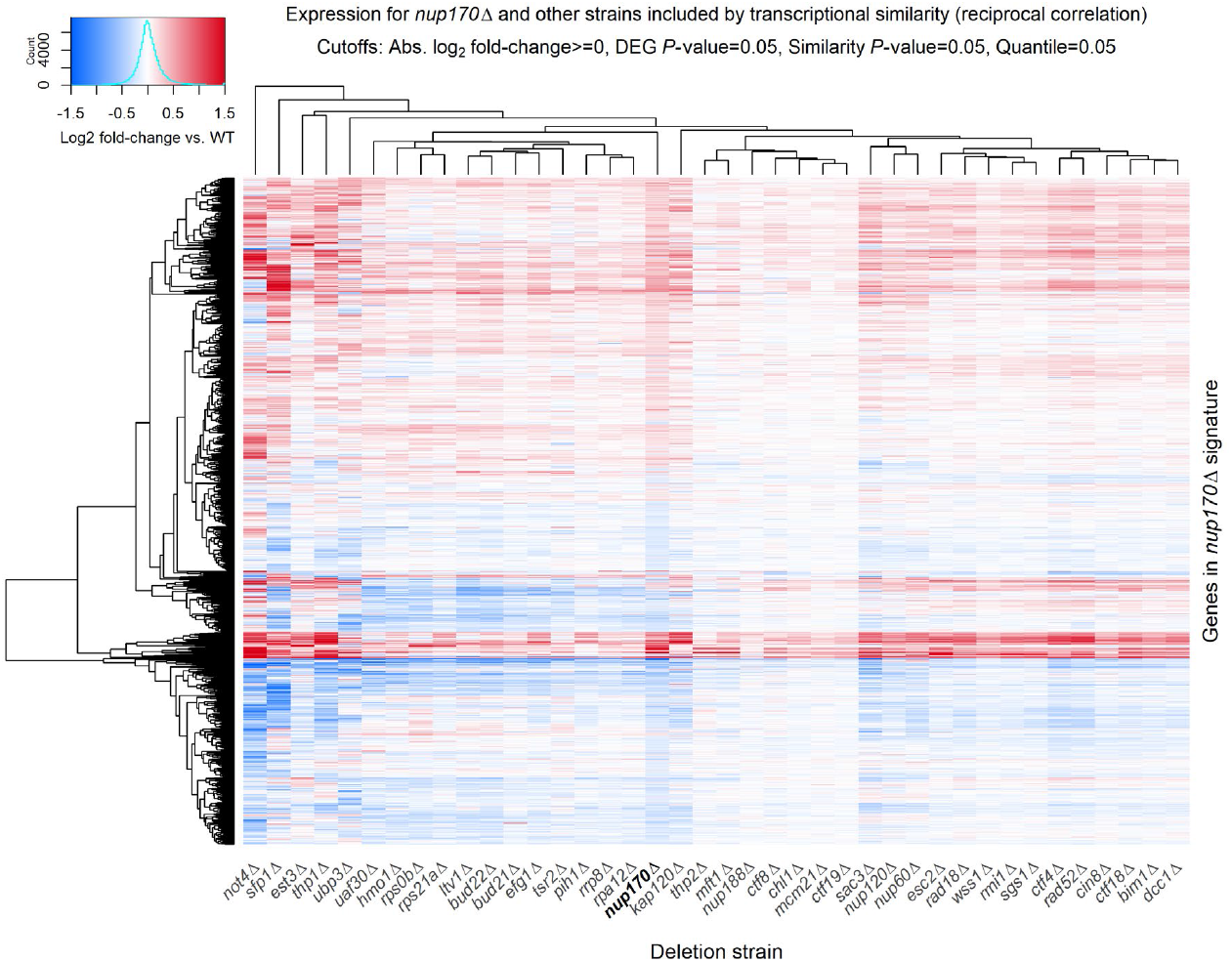
Example visualization provided by DeleteomeTools: a heatmap of log_2_ fold-changes in gene expression for the *nup170*Δ Deleteome strain (bold column label) and 39 other strains with similar transcriptional signatures.

For investigating the genomic position of a transcriptional signature, we provide functions for visualizing the positions of a deletion strain’s signature genes along chromosomes, e.g., from telomeric ends or from the centromere. We have used these visualizations, which were originally devised during early investigations into Nup170’s subtelomeric silencing role (3) and which we refer to as “mountain lake” plots, to identify and compare subtelomeric silencing defects among various Deleteome strains. Supplementary Figure S1 shows an example of this plot for the *nup170*Δ strain.

### Other analysis features

Our codebase also provides a convenience function for performing Gene Ontology (GO) (6, 7) enrichment analyses on an input list of genes that have corresponding deletion strains in the Deleteome. This feature allows users to profile the biological roles of genes deleted in those strains found to be similar to a query strain (see bottom of Figure 1) and thereby predict biological functions for the query gene. If GO enrichment tests on the input gene list show enrichment for a particular biological process, component, or function, that suggests the gene deleted in the query gene may also have a role in that process, function, or component. For example, as shown in Figure 2, we identified 39 deletions strains with transcriptomes that were similar to that of the *nup170*Δ strain. We input the list of 39 genes that were deleted in those strains into our GO enrichment function and the results included significant enrichment for the Ctf18-RFC complex. This is because each of the three Deleteome strains where members of that complex were deleted - the *ctf18*Δ, *ctf8*Δ, and *dcc1*Δ strains - were present in the set of 39 strains classified as transcriptionally similar to the *nup170*Δ strain. This suggested a functional link between Nup170 and the Ctf18-RFC complex. Generally speaking, once a user has identified a set of Deleteome strains of interest (e.g., because they are transcriptionally similar to a query strain) the GO enrichment function allows them to identify significantly enriched biological features among the set of genes that were deleted in those deletion strains. The function relies on enrichment tests provided by the *clusterProfiler* R package (8) and uses the full set of genes deleted in Deleteome strains as the global background for the tests. These enrichment tests account for the varying frequency with which certain GO terms are used as gene annotations reflecting differences in how well certain biological phenomena are studied. Thus, for a frequently applied GO term, a proportionally higher fraction of genes in the test set must share that annotation to yield a significant P-value.

Deleteome Tools also provides convenience functions that quantify the over-representation of genes within subtelomeric or centromeric regions. Given an input set of genes, these functions apply hypergeometric tests to detect enrichment of genes within the specified region. We have used these functions to identify and quantify subtelomeric silencing defects in deletion strains. For example, in the *nup170*Δ strain these tests show a highly significant enrichment of upregulated genes within 25 kilobases (kb) of the telomere (hypergeometric test *P*-value = 3.54e-11). This silencing defect is reflected by the peaks in Figure S1 among upregulated genes located near the telomere.

## Availability and dependencies

DeleteomeTools is implemented as an R package and is publicly available at https://github.com/AitchisonLab/DeleteomeTools.

Identifying similar deletion strains with DeleteomeTools does not require the installation of additional R packages. However, generating visualizations requires the *gplots* and *ggplot2* packages, and performing GO enrichment tests requires the *org*.*Sc*.*sgd*.*db* and *clusterProfiler* R packages, all of which are free and publicly available.

### Threshold optimization and systematic prediction evaluation

Our tool relies on several statistical significance cutoffs to classify a deletion strain as similar to a query strain and then to predict the query strain’s biological function. To find optimal combinations of these cutoffs that maximize predictive sensitivity while limiting the total number of predictions (and thus potential false positives), we performed a systematic analysis where we varied the statistical cutoffs used when a) down-selecting the list of deletion strains with expression signatures similar to a query strain, and b) selecting statistically-significant predicted GO terms from genes deleted in those similar strains. For (a), we varied the percentile threshold to use when selecting similar strains based on reciprocal correlation FDR-adjusted *P*-values. We tested the 99^th^, 95^th^, 90^th^, 80^th^, 50^th^ percentile thresholds and tested omitting the threshold. For (b), we varied the FDR-adjusted *P*-value cutoff to use for selecting enriched GO terms. Values tested were 0.01, 0.05, 0.1, 0.2, 0.5 and 1.0. For each combination of these cutoffs, we predicted GO term annotations for each gene deleted in the Deleteome, and then investigated how well the predicted annotations matched the gene’s reported GO annotations. To assess prediction performance, we focused on how well the tool predicted reported annotations as well as the trade-off between detecting reported annotations and the total number of predicted annotations.

For a binary classification analysis, one could consider the reported GO annotations on a given gene as the “actual positives” and the predicted GO annotations as the “test positives”. Accordingly, an exact match between a predicted and reported annotation would be considered a true positive, and reported annotations that were not predicted would be considered false negatives. However, we considered the set of “actual negative” conditions (the GO annotations *not* applicable to a gene) as unknown. Assuming otherwise presumes that the functions of all genes analyzed here have been fully characterized and the GO annotations associated with each gene can be considered a complete account of their functions. As exemplified by our discovery of the novel relationship between NUP170 and the PCNA machinery, we know this is not the case, as the existing GO annotations on NUP170 do not capture this relationship. Thus, for our analysis, we did not include predictive measures such as specificity or recall that rely on true negative or false positive quantification.

Furthermore, rather than classify a predicted GO annotation in a binary fashion based on whether it was also a reported annotation, we aimed to quantify how *similar* each predicted annotation was to a reported annotation. Even though a predicted GO annotation might not match a reported annotation exactly, it may be very closely related to it, and therefore still capture an informative aspect of the gene’s functionality. Thus, we quantified the similarity between predicted and reported GO annotations using semantic similarity metrics. A variety of these metrics exist (9), and we used three of the six that are available in the *GOSemSim* R package (10) (version 2.30.2). We required that the metric quantify the similarity between two identical GO term IDs as 1.0 and this was the case for the “Lin” (12), “Jiang” (13), and “Wang” (14) metrics. The first two approaches are information content-based, and the third is graph-based.

For each combination of statistical cutoffs we tested, we predicted GO annotations for every gene represented by a deletion strain in the Deleteome using the reciprocal correlation approach. We then compared predicted to reported annotations, recording the semantic similarity score between each reported annotation and its closest match among the predicted annotations. Next, we computed the median of these “best match” scores across all Deleteome strains as well as the median ratio of number of predicted annotations versus reported annotations. We then identified the optimal combination of statistical cutoffs by determining which pair of these two medians was closest to the point of best possible performance (which is where the median semantic similarity of best-matching annotations and the ratio of the number of predicted annotations to reported annotations are both 1.0). To determine whether optimal cutoffs would be consistent across annotations from the three GO sub-ontologies (cellular component, biological process, and molecular function), we examined prediction performance separately across each sub-ontology. Using the optimal set of statistical cutoffs identified, we then examined the overall predictive performance of our tool by quantifying the fraction of all reported GO:*biological process* annotations that were detected by our approach at varying levels of semantic similarity. To compare the performance of our tool against results obtained by chance, we performed this same systematic evaluation but instead of using the reciprocal correlation method to identify deletion strains similar to a query strain, the similar strains were selected at random. For each query strain in this comparison method, the number of randomly-selected similar strains was the same as the number identified by the reciprocal correlation method.

In line with the Critical Assessment of Functional Annotation (CAFA) challenge (11) – an ongoing competition to accurately predict GO annotations for proteins – our evaluations only used reported GO annotations with evidence codes indicating experimental or high-throughput evidence, a traceable author statement, or curator inference. Reported GO annotations were obtained from the *org*.*Sc*.*sgd*.*db* R package (version 3.19.1).

## RESULTS

To predict a gene’s biological function using DeleteomeTools, users enter the gene name into a package function that 1) retrieves the transcriptome data for the Deleteome strain in which the gene was deleted, 2) iteratively compares that transcriptome to all other Deleteome strain transcriptomes, and then 3) lists the genes that were deleted in strains that had similar transcriptomes. The resulting gene list includes candidates that are predicted to functionally interact with the input “query” gene because their deletion caused similar transcriptomic changes. Users can then use DeleteomeTools to predict the query gene’s biological functions by performing GO enrichment analysis on the list of genes generated in step 3. If the analysis identifies significant enrichment for a specific GO:*biological process*, GO:*cellular component*, or GO:*molecular function* annotation, it suggests that the query gene may be associated with that biological role.

As detailed in Materials and Methods, DeleteomeTools offers two complementary methods for the similarity analysis in steps 1-3. Both methods begin by identifying the differentially expressed genes in the strain where the query gene was deleted (the strain’s “signature”). The first approach quantifies similarity by performing correlation tests between strain signatures and the second uses hypergeometric enrichment tests to assess qualitative overlap between signatures. Both similarity tests, as well as the GO enrichment tests for functional prediction, use numerical thresholds to distinguish significant from non-significant results, and adjusting these cutoffs impacts the sensitivity of our tool’s predictions. Thus, we aimed to find an optimal combination of statistical thresholds that would maximize prediction sensitivity while minimizing the total number of predictions. Here we report the results of these analyses where we tested various combinations of two key thresholds used in our pipeline: 1) the percentile used to down-select strains similar to a query strain and 2) the FDR-adjusted *P*-value cutoff for GO enrichment tests. We sought to find a combination that best predicted previously reported GO annotations for query genes. Using these optimized thresholds, we then performed a systematic evaluation of our tool’s predictive capabilities by predicting functions for all genes with deletion strains in the Deleteome (i.e., any gene that can be a “query” gene) and comparing them to known gene functions. For this assessment, we compared the predicted GO annotations for the gene to its reported GO annotations. As described in Materials and Methods, we used semantic similarity metrics for this comparison to quantify the degree of functional agreement as a continuous measure rather than a binary match/no-match outcome.

### Optimized threshold values

For GO:*cellular component* annotations, using the 99^th^ percentile for strain similarity and 0.2 for the GO enrichment significance cutoff gave the best performance in two out of three semantic similarity measures. These same cutoffs gave the best performance for GO:*molecular function* annotations across all three measures. For GO:*biological process* annotations, using the 90th percentile for strain-similarity and 0.1 for the GO enrichment significance cutoff gave the best performance in all three measures (Figure S2). Our results showed that the optimal statistical thresholds depend on the GO sub-ontology containing the annotations. DeleteomeTools users may wish to adjust their thresholds depending on which type of GO annotation they consider most critical to their analyses.

### Predictive performance

Using the optimal thresholds for detecting GO:*biological process* annotations reported above, we predicted GO annotations for each gene deleted in the Deleteome strains. We then used semantic similarity to identify the best-matching predicted GO annotation for each reported annotation on those genes. Out of 1,474 Deleteome query strains, 1,341 had at least 3 differentially-expressed genes - the technical minimum required for our correlation-based analyses. Among these, 1,109 had at least one similar deletion strain based on our reciprocal-correlation approach. From this set, 1,050 had at least one reported GO:*biological process* annotation for their deleted gene. For 846 of these genes, the tool predicted one or more GO:*biological process* annotations, and we assessed the predictive performance of our tool using the reported annotations from this gene subset. Table S1 lists all 1,474 Deleteome query strains and indicates which were included in our predictive performance evaluation.

Median semantic similarity of the best-matching predicted annotation was 0.72, 0.71, and 0.63 for the Jiang, Lin, and Wang semantic similarity metrics, respectively (Figure 3A). While these results vary according to semantic similarity metric, they consistently show that median scores from the reciprocal correlation method are significantly higher than scores resulting from randomly selecting similar strains for a query strain (Mann-Whitney U test *P*-values ≤ 3.47e-45). Given that several of our tool’s functional predictions for NUP170 were ultimately validated experimentally, we investigated whether the tool’s performance for NUP170’s annotations was substantially better than its overall performance across all annotations, thereby leading to more accurate predictions for that gene. However, we found this was not the case. The median semantic similarity scores for NUP170’s best-matching predictions were 0.81, 0.75, and 0.55 for the Jiang, Lin, and Wang metrics, respectively. These values correspond to the 61^st^, 55^th^, and 42^nd^ percentiles of all semantic similarity scores, indicating that the tool’s performance for NUP170 functional prediction was not an outlier.

**Figure 3.**
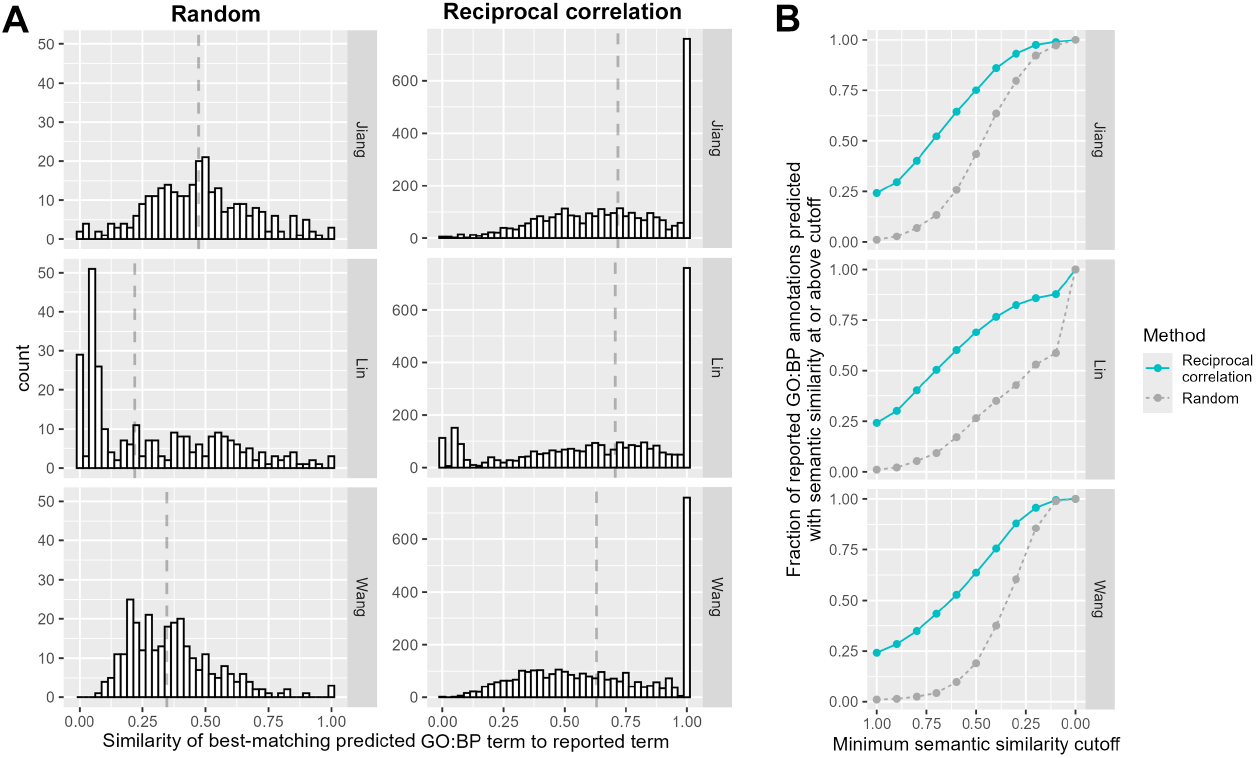
Overall performance of DeleteomeTools for predicting reported GO:*biological process* (GO:BP) annotations. **A**) Semantic similarity distributions for best matches between reported and predicted annotations using reciprocal correlation (right) or random (left) approach. Dashed vertical lines indicate distribution medians. For each semantic similarity method (rows), the median value of the reciprocal correlation method is significantly higher than the random method (Mann-Whitney U test P-values ≤ 3.47e-45). **B**) Performance quantified at various levels of semantic similarity across similarity metrics. X-axis: The minimum semantic similarity between a predicted and reported annotation for a prediction to count. Y-axis: the fraction of reported GO:BP annotations predicted by reciprocal correlation (solid, blue line) or randomly (dashed, gray line) with a semantic similarity score at or above the cutoff. Results for A and B generated using optimal statistical cutoffs for the GO:BP process sub-ontology as described in Material and Methods and Results.

From our full set of evaluation results, we quantified the probability of a reported GO:*biological process* annotation being predicted with at least certain minimum semantic similarity (Figure 3B). For example, by the “Lin” metric, there is a 24% chance of a reported GO annotation being matched exactly by a predicted annotation (i.e., with a semantic similarity of 1.0). By the same metric, there is a 50% chance of matching a reported annotation with a similarity ≥ 0.7 and a 69% chance of matching with a similarity ≥ 0.5. To illustrate the degree of relatedness between GO terms with similarity scores of 0.7 and 0.5, we provide examples of predicted annotations for Nup170 that matched reported annotations at those levels (Table S2).

## DISCUSSION

Our performance analysis indicates that our tool predicts reported GO:*biological process* annotations with far greater frequency than expected by chance and that a majority of reported annotations on the 846 genes we tested were predicted with moderate or stronger semantic similarity. These results are based on analyses that used statistical cutoffs optimized to balance reported annotation detection with total number of predicted annotations. Users may wish to adjust these cutoffs to favor the former or latter depending on their analysis needs.

A distinguishing feature of our approach for quantifying similarity between deletion strains is that we limit our strain-to-strain comparisons to gene expression values that changed significantly versus wild-type. This contrasts with other approaches that cluster deletion strains based on the full complement of measured gene expression values. This may have important implications for predicting gene function. For example, whereas our similarity analysis predicted a functional relationship between Nup170 and members of the Ctf18-RFC complex with high confidence, a UMAP-based clustering of the Deleteome placed Nup170 and the Ctf18-RFC members in distinct clusters that are relatively distant from each other in UMAP space (2). We suspect this may be due to differences in the complement of gene expression values used to quantify strain similarity. Within this UMAP cluster, Nup170 is grouped with 18 other members, and together they circumscribe a set of biological functions distinct from those of the Ctf18-RFC complex. Ten of these 18 members are also predicted to be co-functional with Nup170 by our reciprocal correlation approach. These findings suggest that while there may be substantial agreement between functional predictions made using full transcriptomes and signature-based approaches such as ours, the codebase described here offers a valuable, complementary strategy for prediction that may highlight important functional relationships between genes not revealed by other methods. We note that application of these signature-based approaches is not limited to the Deleteome dataset but could potentially be applied to predict gene function in other organisms where sufficient transcriptome profiles associated with gene disruptions are available for analysis.

## DATA AVAILABILITY

The DeleteomeTools package, which includes all data used in our analyses, is publicly available at https://github.com/AitchisonLab/DeleteomeTools.

## ACKNOWLEDGEMENTS

We thank Drs. Fred Mast, Fergal Duffy, and Santhosh Sariki for their help testing DeleteomeTools. We also thank Dr. Marc Carlson for technical guidance on R packaging.

