## Supplementary for "DeleteomeTools: Utilizing a compendium of yeast deletion strain transcriptomes to identify co-functional genes": SupplementaryFigures.docx

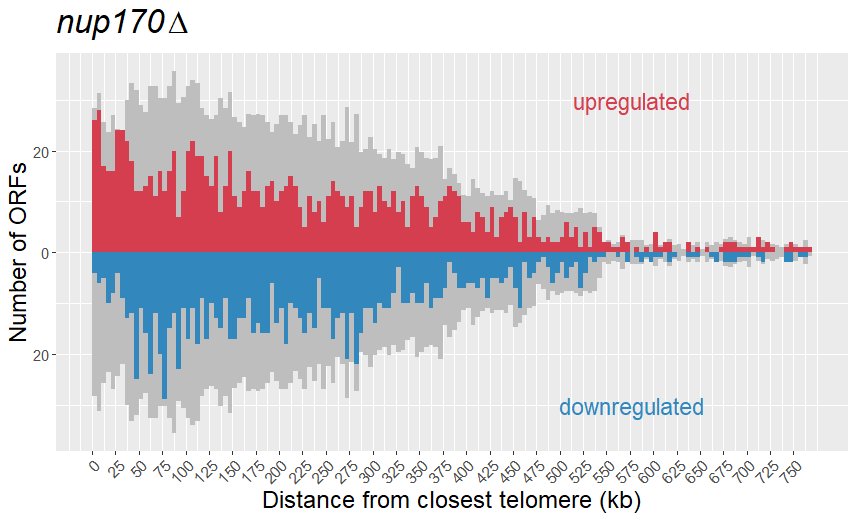

**Figure S1.** **Visualizing the genomic position of differentially-expressed genes in the *nup170*Δ deletion strain from the Deleteome.** Red indicates significantly upregulated open reading frames (ORFs), blue indicates significantly downregulated ORFs. Gray indicates the background distribution of all ORFs in the yeast genome where, for visualization purposes, the number of ORFs shown is a third of the total in each x-axis bin. This background distribution is mirrored above and below the x-axis.

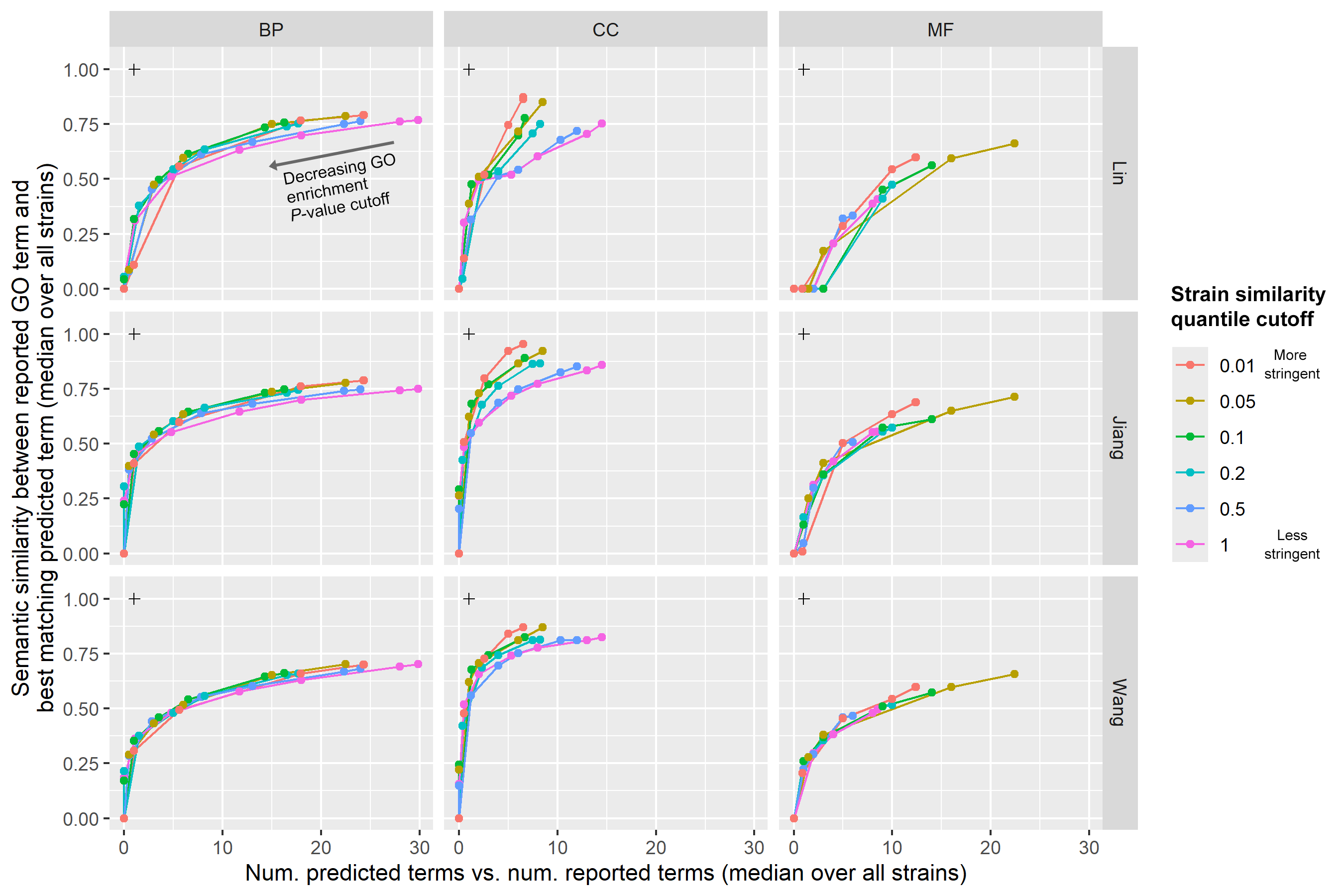

**Figure S2.** **Identifying optimal parameter threshold values for predicting GO annotations on genes deleted in the Deleteome.** We evaluated GO annotation predictions using various combinations of two thresholding parameters, the first of which determines the number of genes qualifying as similar to a query gene (the “Strain similarity quantile cutoff”) and the second which is the FDR-adjusted *P*-value significance cutoff for GO enrichment tests on those genes. For each line in each plot, the GO enrichment cutoff decreases from right to left. The “+” symbol indicates the point of optimal performance, and the best-performing combinations of parameter values were identified as those giving prediction results closest to this point. Columns show performance for annotations from the three different GO sub-ontologies: biological process (BP), cellular component (CC) and molecular function (MF). Rows indicate performance for the three different semantic similarity metrics tested.
